# Beyond DNA barcodes: an open-source workflow for recovering and organizing barcoded vouchers for ecological and evolutionary research

**DOI:** 10.64898/2026.08.06.743289

**Authors:** Vivian Feng, Huei-Min Lin, Amrita Srivathsan, Hui Wang, Leshon Lee, Ronniel Pedales, Dirk Oberschmidt, Rudolf Meier

## Abstract

1. Most species are neither discovered nor named, let alone included in analyses that require biological information such as trait measurements, images, ecological information and genome-scale data. Specimen-level DNA barcoding can help discover many of these species rapidly, but everything beyond discovery requires vouchers organized into putative species. Yet, existing barcoding workflows lack efficient techniques for voucher recovery, creating a post-barcoding bottleneck that limits the ability of converting barcoded specimens into biological knowledge.

2. Here we present a low-cost, open-source workflow consisting of two stages. The first safeguards barcoded specimens by separating them from DNA extracts and transferring them from microplates into ethanol-filled glass vials. The second converts the resulting voucher collection into a searchable physical resource by linking barcode-derived molecular Operational Taxonomic Unit (mOTU) assignments to vial positions and enabling specimens to be sorted into putative species either manually or automatically using a newly developed open-access robot (*SORTER*).

3. We evaluated the workflow using 2,024 insect specimens distributed across 21 96-well plates. For the first stage, DNA separation and specimen transfer required approximately 15 minutes per plate. For the second stage, *MOTUmapper* generated retrieval coordinates in a few seconds, after which the 2,024 vouchers belonging to the 452 putative species could be recovered manually in 5 days or with *SORTER* in 5 hours. Throughout both stages, specimen identities remained linked to barcode sequences, metadata and storage positions.

4. Vouchers are the Rosetta stones of biology because they connect different kinds of data to the same specimens. By safeguarding these vouchers and making them searchable, the workflow converts barcode projects from one-time molecular surveys into reusable resources for ecological and evolutionary research.

## 1 Introduction

Specimen-level DNA barcoding is increasingly used to generate large biodiversity datasets for poorly known and hyperdiverse taxa, routinely encompassing many species that remain undiscovered or unnamed (Mora et al., 2011; Hortal et al., 2015; Srivathsan et al., 2019; Meier et al., 2025a; Meier et al., 2025b; Colwell et al., 2026). For example, the Barcode of Life Data Systems alone now contains more than 20 million barcode records for the kingdom Animalia, with only 6.6 million records identified to species (accessed 04 August 2026; Ratnasingham et al., 2024). These barcoded specimens are valuable beyond yielding a barcode as they are associated with metadata such as collection locality and date. They can support holistic insect monitoring (Meier et al., 2024), genomic applications (Feng et al., 2026), trait-based analyses (Shirali et al., 2026), and images for the training of machine-learning systems (Gharaee et al., 2025). All these downstream uses, however, depend on access to the barcoded specimens (Goldblatt et al., 1992; Funk et al., 2005; Troudet et al., 2018) that are the primary evidence linking sequence data, metadata and organismal traits (Huber, 1998; Ruedas et al., 2000; Suarez & Tsutsui, 2004).

Unlocking the full potential of specimen-level barcode datasets thus requires workflows that keep specimens safe and accessible after sequencing. Keeping specimens accessible is becoming a serious bottleneck because a single bulk sample of invertebrates can yield thousands of specimens distributed across hundreds of 96-well microplates representing hundreds of species (Srivathsan et al., 2019). Currently, the only option for recovering these specimens is using forceps to manually move them into vials. However, this is slow, repetitive and difficult to scale without errors. The alternative is leaving the specimens in 96-well microplates or other molecular-laboratory consumables. However, here the evaporation of preservatives and plastic degradation threatens the long-term survival of the specimens. These means that they are just “implicit vouchers” with the associated data being at risk of becoming “ghost records” (Portman et al., 2025). Addressing this bottleneck requires two linked processes. The first is safeguarding vouchers by transferring them into durable storage while preserving their association with barcode records and metadata. The second stage is to make those vouchers findable by reorganizing them by putative species. This is a non-trivial task because specimens belonging to the same molecular operational taxonomic unit (mOTU) may be scattered across many plates and wells. Here, we present low-cost, open-source workflows addressing both stages of voucher recovery. The first stage safeguards barcoded specimens by separating them from their DNA extracts and transferring them from microplates into preservative-filled glass vials while maintaining their positional identities. The second stage links barcode-derived mOTU assignments to vial positions, allowing specimens belonging to the same putative species to be retrieved manually or reorganized automatically with a newly developed and open access *SORTER* robot. We evaluate the workflow using 2,024 insect specimens and quantify processing time, retrieval time and implementation costs. Together, the workflow converts barcoded specimens into durable, searchable resources for taxonomy, trait research, imaging and genomics.

## 2 Materials and Methods

### Stage 1: Safeguarding vouchers

#### 2.1 The inner-outer plate system

The central challenge in preserving vouchers during DNA barcoding is exchanging liquids around fragile specimens at 96-well scale without causing damage: specimens from a bulk sample must first be individualized in wells, the transfer liquid removed, DNA extraction buffer added, and the DNA extract recovered before each voucher can be transferred into a vial for long-term storage. We would argue that this is best achieved with a sieve that keeps the specimen safe while allowing the surrounding liquids to be exchanged.

In our workflow, we use the wells of an “inner plate” as a sieve. A small hole at the base of each well retains liquid through surface tension but allows it to be passed through the opening when centrifuged (Fig. 1). Once specimens have been placed into the wells of an inner plate, liquids can be exchanged by placing this plate above a regular “outer plate” and then pushing the liquid from the inner to the outer plate using a plate centrifuge. Inner plates with holes in their wells can be produced easily from standard 96-well plates (e.g., catalogue no. 211-0262, VWR, Pennsylvania, USA) by piercing the base of each well with a size 1 insect pin (e.g., Entomoravia, Slavkov u Brna, Czech Republic; time needed: 5 min) or one can use a CO₂ laser engraver for fast production of many plates (see Supplementary Material for printing file and instructions; time needed: 30 s). For centrifugation any plate centrifuge can be used that allows for two stacked plates to be spun at 1,000 rpm for 1 min (e.g. Sorvall X4 Pro: Thermo Fisher Scientific, Massachusetts, USA).

**Figure 1.**
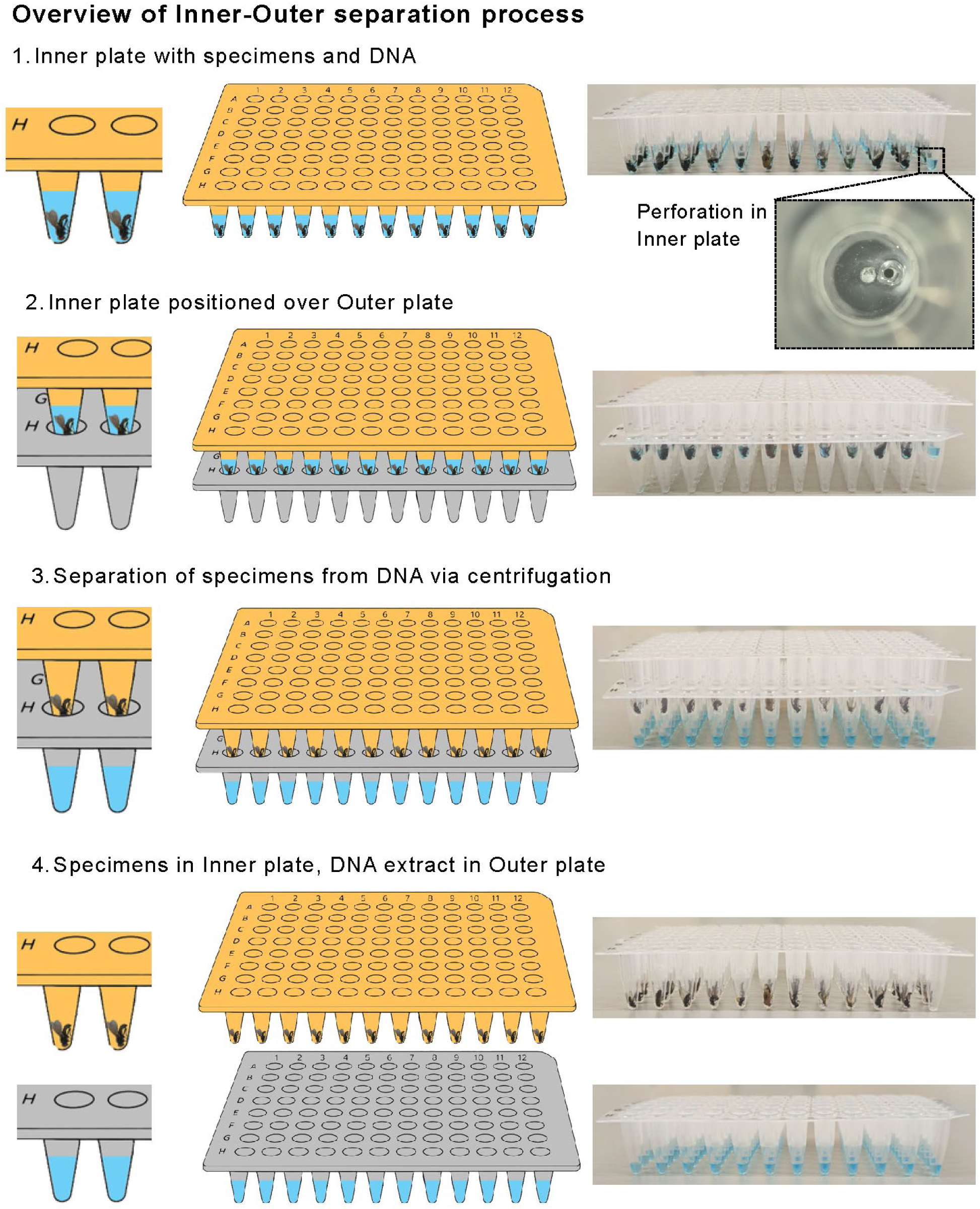
Inner-Outer separation system. Specimens are prepared with DNA extraction buffer in the Inner plate (yellow), which is perforated at the bottom of each well, as shown in the insert. Following DNA extraction, the inner plate is positioned above the outer plate (grey). Centrifugation then separates the specimens from the DNA extract (blue).

#### 2.2 Transfer of specimens into glass vials

After removal of the DNA extract into the outer plate, 95 specimens remain in 95 wells of the inner plate (we use one well of the 96-well plate as negative control). All 95 specimens can be transferred simultaneously into 95 storage vials using a “popsicle” device (Fig. 2). The device consists of a 3D-printed holder carrying 96 screws, each 1.6 mm in diameter and 16 mm long (article no. 59600070009-1000, Schraubenluchs, Marienberg, Germany). The spacing of the screws is aligned with the well spacing of a 96-well plate. Depending on specimen size, 100–120 µl of water is added to each occupied well before the popsicle device is inserted (see Fig. 2). The plate-popsicle sandwich is then frozen at −20 °C for at least 6 h. Freezing creates 95 icicles that enclose the specimens. To remove the frozen specimens without breaking the icicles, the plate is briefly immersed in tap water to release the frozen plugs. When the device is then lifted, all specimen-icicles are freed from the inner plate and can be lowered into a 3D-printed rack containing 95 labelled glass vials (e.g., article no. 70202.1, Macherey-Nagel, Düren, Germany). The spacing of the vials again matches the spacing of the 96-well plate. Each vial is filled with 800 µl of 96% ethanol so that the final storage concentration of the ethanol is approximately 80% after the icicles melt and release the specimens into the vials. Afterwards, the vials can be sealed with caps (e.g., article no. 702807, Macherey-Nagel, Düren, Germany).

**Figure 2.**
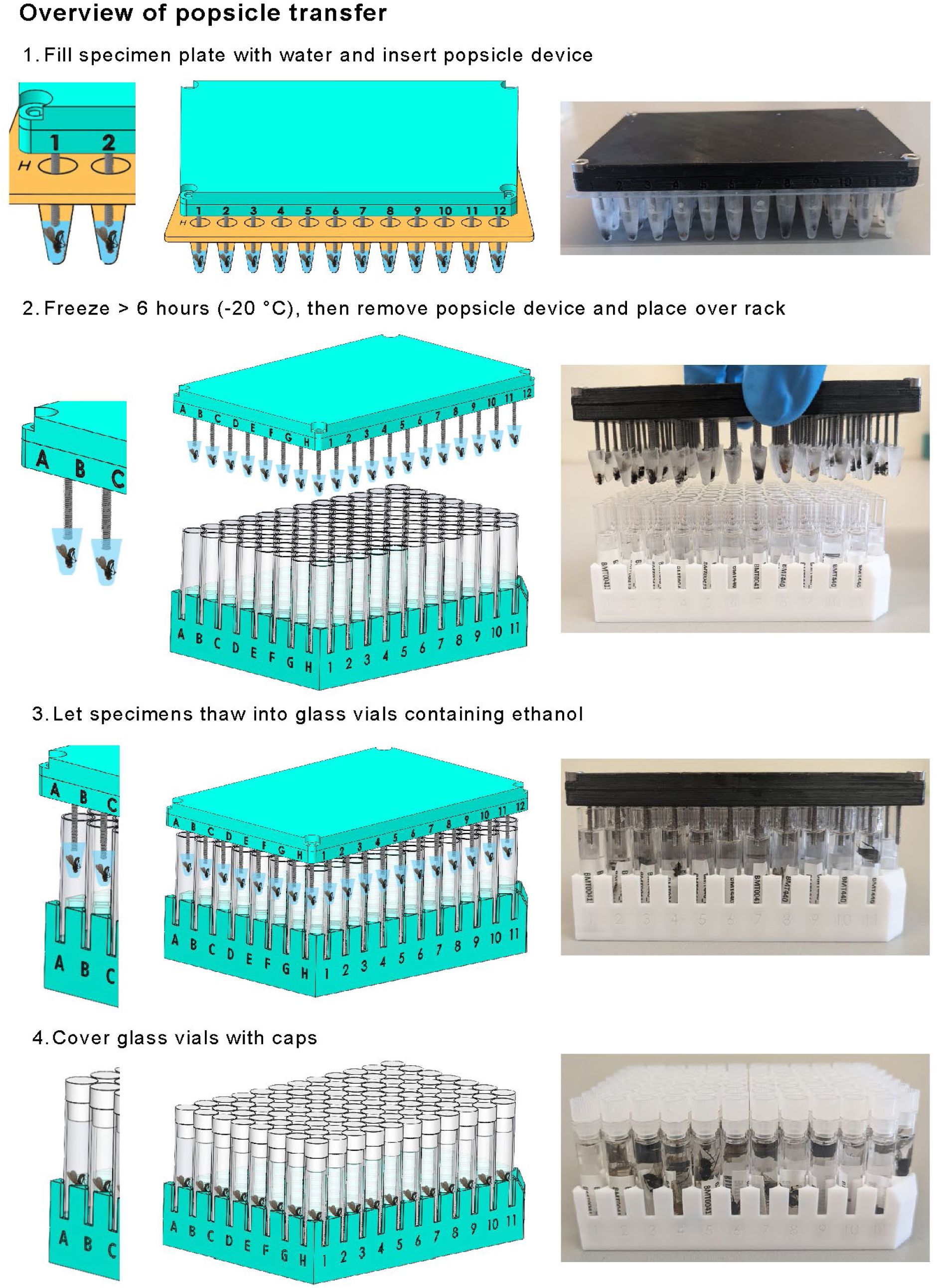
The popsicle transfer workflow.

### Stage 2: Making vouchers findable

#### 2.3 Linking barcodes, mOTUs and vial positions

To retrieve vouchers belonging to the same mOTU, each physical specimen must remain linked to its barcode sequence, which can then be used to assign the specimen to a mOTU. In tagged amplicon sequencing, the first task is accomplished by using a unique combination of sequence tags incorporated into the forward and reverse primers of each specimen (Meier et al., 2016; Srivathsan et al., 2019, 2021; Hebert et al., 2025). A demultiplexing file records the correspondence between these tag combinations and specimen location in plates and wells. The second task is using barcodes to assign each specimen to a mOTU. There are many different clustering and mOTU-delimitation methods and recent efforts have led to formulation of a standardized export format (SPART: Species PARTition; Miralles et al., 2022). SPART outputs can be obtained easily for ASAP (Puillandre et al., 2021), ABGD (Puillandre et al., 2012) and Objective Clustering (Meier et al., 2006; Srivathsan et al., 2025), and viewed online (Spart Explorer: spartexplorer.mnhn.fr) or locally (Srivathsan et al., 2025). We here introduce a new software package *MOTUmapper* that uses the SPART and demultiplexing files to link barcodes to mOTUs and mOTUs to vial positions. *MOTUmapper* generates a CSV sorting sheet that groups vouchers by mOTU and reports the physical location of every specimen assigned to each mOTU. *MOTUmapper* includes features to produce the demultiplexing file as well and is a light-weight tool written in HTML and JavaScript, which avoids software installations. It brings together well-tested Python-based software that were converted into a simplified web design that was facilitated using large language models (LLMs). The authors thus validated the *MOTUmapper* by comparing results to output from legacy Python scripts.

#### 2.4 Manual and robotic specimen sorting

The sorting sheet generated by *MOTUmapper* can be used directly to retrieve vouchers manually from the vial racks (Fig. 3). This is suitable when only a few hundred vouchers or mOTUs need to be retrieved. For larger datasets, a complete reorganization of the collection is desirable. This can also be accomplished manually but is time-consuming. Therefore, we also developed a new *SORTER* robot that automatically sorts the vials by mOTU. As with manual sorting, *SORTER* sorts one mOTU after another. Specimens without a barcode are grouped by *MOTUmapper* and processed in the same manner as a mOTU either manually or with *SORTER*.

**Figure 3.**
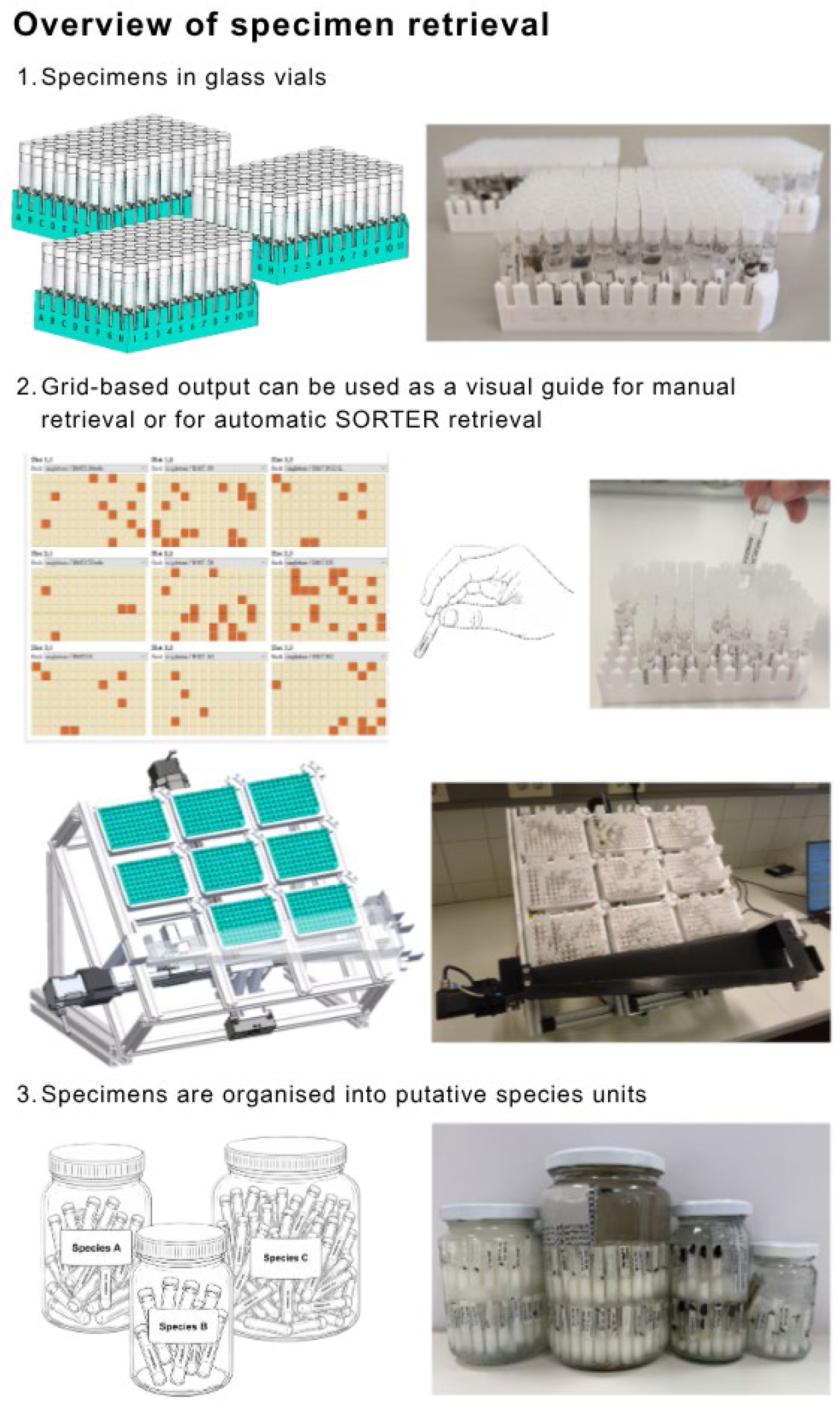
Locating priority taxa and sorting individuals to putative species units.]

The current prototype accommodates approximately 850 vials (Fig. 3). Larger collections therefore require rack changes or several *SORTER* units to operate in parallel. A video of the *SORTER* in action is available at https://youtu.be/0p4tw--LtPc.

The nine racks of the *SORTER* are mounted at an adjustable incline so that vials displaced from their positions slide into a collection chute so that the mOTUs can be “harvested”. Vials are ejected by a replaceable 3D-printed pin moved along the x-, y- and z-axes by stepper motors. Limit switches define the home position of each axis during initialization. *SORTER* is controlled through a Python-based graphical user interface (GUI) that loads the sorting sheet, assigns rack positions, selects vials and executes the retrieval sequence. LLMs were used solely to assist with modifications to the GUI and were not used to develop or alter the underlying control logic or functionality. Machine-specific parameters, including calibration coordinates, vial spacing and movement timing, are stored in a TOML configuration file. The Python program communicates with an Arduino controlling the motors through a USB serial connection. Retrieval order is optimized using a nearest-neighbour algorithm based on Manhattan distance, *D* =∣ *x*_1_ − *x*_2_ ∣ +∣ *y*_1_ − *y*_2_ ∣, which directs the movement of the device along the x- and y-axes.

#### 2.5 Workflow evaluation

We evaluated the workflow using 2,024 insect specimens distributed across 21 plates. Cytochrome oxidase I (COI) barcodes were generated using the protocol outlined in Srivathsan et al. (2024), and sequences passing the quality check were clustered into mOTUs at 3% uncorrected p-distance using *IntegraTax* (Srivathsan et al., 2025). The resulting partitions were exported in SPART format and linked to vial positions with *MOTUmapper*. For each workflow component, we recorded active processing time separately from unattended steps such as freezing and 3D printing. Per-plate times were measured for Inner–Outer separation, popsicle transfer, preparation of the vial racks and closure of the vials. *MOTUmapper* runtime was measured for the complete dataset. Retrieval of all vouchers was performed both manually and with *SORTER*, and total retrieval time was recorded for each approach. Implementation costs were calculated separately for reusable equipment and recurring consumables.

## 3 Results

### 3.1 Pipeline throughput and efficiency

When applied to 2,024 insect specimens distributed across 21 plates, we found that 1,583 specimens yielded COI barcodes that belonged to 452 mOTUs based on an uncorrected 3% p-distance threshold (Srivathsan et al., 2025). Most steps of the workflow only required minutes of active handling per plate (see Table 1), while freezing accounted for most of the elapsed time but required no operator involvement. *MOTUmapper* generated the retrieval file for the complete dataset in <30 s. Reorganization of all vouchers required approximately five working days manually, compared with approximately 5 h using *SORTER*.

**Table 1.**
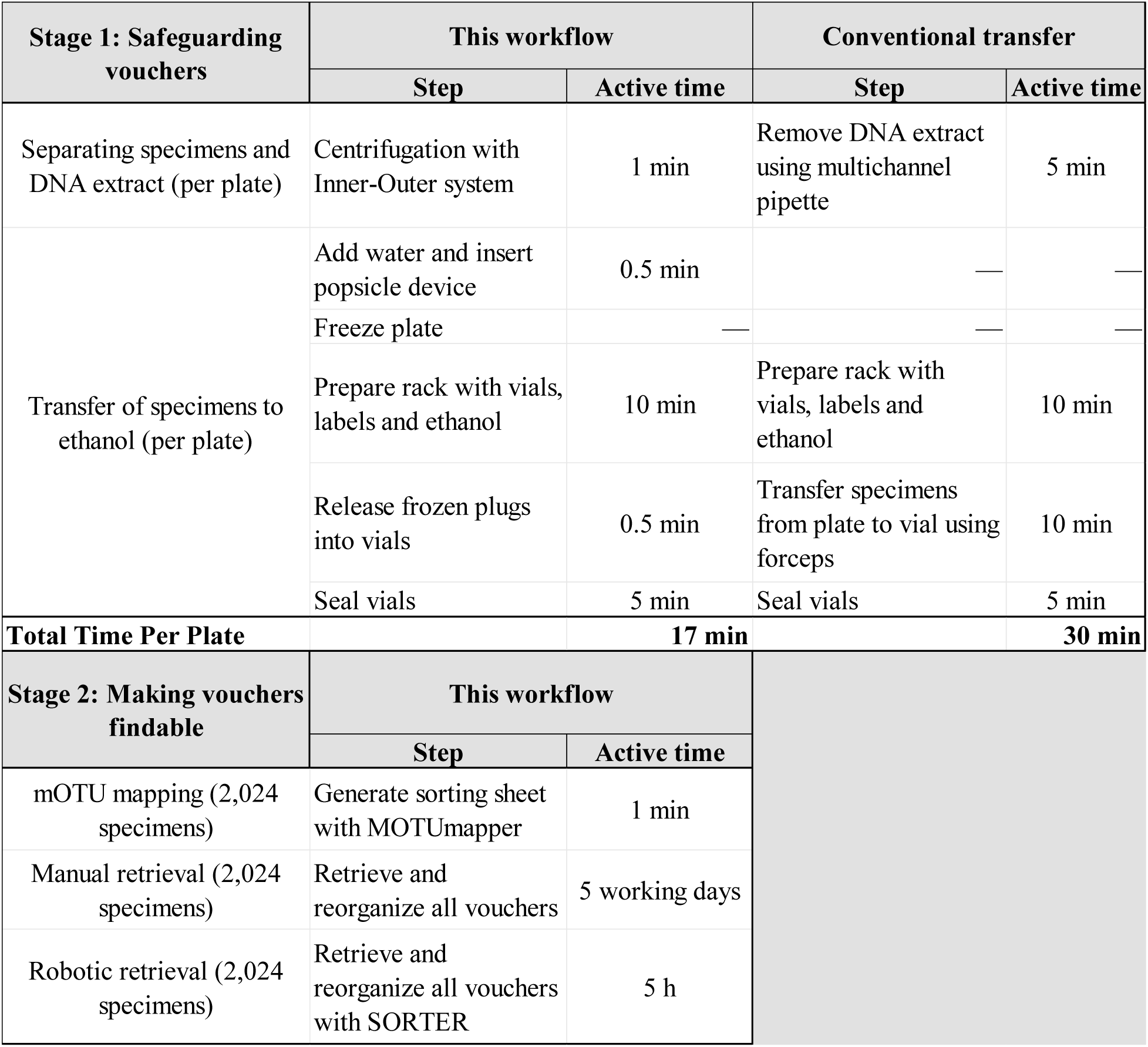
Steps and active time requirements for the two-stage voucher recovery workflow. For Stage 1, freezing time is excluded and the workflow is compared with conventional manual transfer of individual specimens. Reported values are mean timings from three students performing each procedure.

Preparation of the reusable equipment was a one-time investment. An inner plate could be produced manually in approximately 5 min and with a laser engraver in 30 s. Printing the popsicle holder and vial rack required approximately 6 and 12 h, respectively, followed by 10 min to assemble the popsicle device. These printing times were unattended, and the components could subsequently be reused for multiple plates.

### 3.2 Low-cost and open-source implementation

The basic configuration of the workflow uses manually perforated inner plates and manual specimen retrieval and therefore requires no specialized equipment beyond a plate centrifuge and access to a 3D printer. The reusable popsicle device requires 96 screws (ca. €2.50), together with a 3D-printed screw holder (ca. €1.80) and vial rack (ca. €1.20). For higher-throughput applications, inner plates can be perforated with a CO₂ laser engraver and specimens retrieved with *SORTER*. Fabrication of *SORTER* required approximately 130 h of unattended 3D printing and 2.5 days of assembly, with component costs of approximately €1,600. All custom designs, *MOTUmapper* and the *SORTER* control software are open-source, allowing laboratories to reproduce or modify the workflow according to their processing volume and available infrastructure.

## 4 Discussion

Vouchers are the Rosetta stones of biology because they connect data to organisms, species names and the accumulated literature on their biology. Their importance is evident when considering which of Hortal et al.’s (2015) seven shortfalls in biodiversity knowledge can only be addressed with specimens. The dependence on vouchers is strongest for the Linnean shortfall, because taxonomic work requires access specimens and species identifications should be linked to physical references (Wang et al., 2018; Hartop et al., 2022). The scale of this task is illustrated by a recent metabarcoding study of German Malaise-trap samples, which recovered 10,803 validated insect species but estimated a further 21,043 plausible species lacking reference barcodes or names (Buchner et al., 2025). Retrievable specimens could allow many of these anonymous mOTUs to be matched to described species and their published biology. Vouchers are also central to reducing the Darwinian, Raunkiæran and Eltonian shortfalls because they provide material for reconstructing evolutionary relationships, measuring phenotypic traits and documenting biotic associations. By contrast, Hortal et al.’s (2015) Wallacean and Prestonian shortfalls, concerning species distributions and abundances, can increasingly be addressed from molecular data alone, whereas the Hutchinsonian shortfall (data on environmental tolerances) usually requires experiments with living or freshly collected organisms.

The workflow presented here was thus developed to preserve biology’s Rosetta stones and organizing them into units useful for ecological and evolutionary studies. First, the Inner–Outer plate system and popsicle transfer move specimens from microplates unsuitable for long-term storage into durable individual vials without breaking their positional identity. Second, *MOTUmapper* links barcode-derived partitions to the physical locations of the vouchers, allowing specimens to be retrieved manually or reorganized with *SORTER*. Applied to 2,024 specimens, the workflow linked 1,583 barcodes representing 452 mOTUs to their storage positions. The complete dataset was mapped in under 30 s, and retrieval was reduced from about five working days manually to approximately five hours with *SORTER*.

The first stage of the workflow addresses the material legacy of DNA barcoding studies. Non-destructive extraction methods developed over the years enable preservation of intact specimens (Castalanelli et al., 2010; Kirse et al., 2023; Santos et al., 2018; Stein et al., 2022). Specimens processed in such a manner are left in extraction plates in theory, but microplates and other molecular consumables were not designed for long-term storage. The Inner–Outer system and popsicle transfer make permanent storage part of the molecular workflow. The main advantage is that liquid exchange or voucher transfer can be performed at plate-scale while retaining the correspondence among specimen, DNA extract, barcode and storage position. All equipment is reusable, inexpensive and can be produced with widely available fabrication methods. These are important criteria because most of the unknown biodiversity resides in countries with limited science funding (Brydegaard et al., 2024).

The second stage of the workflow makes vouchers findable. In non-targeted samples, specimens belonging to the same mOTU may be scattered across many plates, racks and wells. *MOTUmapper* converts a molecular signature into a plate position captured in a retrieval sheet, allowing users to assemble the material required for a question. This could be representatives of common, rare or unusual mOTUs, taxa driving differences among sites, or closely related taxa selected for comparative analysis. Complete reorganization is optional and the user can also opt to keep the original vial arrangement and only use *MOTUmapper* to find specific specimens. This flexibility is strengthened by the use of SPART which allows the user to decide which mOTU clustering algorithm to use.

The workflow presented here is deliberately flexible because methods intended for global biodiversity research must accommodate laboratories with very different budgets, infrastructure and technical capacities (Brydegaard et al., 2024). This follows the same design principles that guided our development of imaging tools ranging from a frugal, do-it-yourself microscope to high-throughput robotic systems (Wührl et al., 2022, 2024). Manual voucher retrieval is sufficient for laboratories that barcode a few hundred or thousand specimens only occasionally, whereas *SORTER* becomes advantageous when thousands of vials must routinely be assembled into hundreds of mOTUs. The other components can likewise be adopted incrementally: Inner plates may be perforated manually or by laser, and robotic retrieval added only when justified by throughput. Open-source hardware and software further enable local fabrication, repair, modification and adaptation to different storage formats.

The workflow has the obvious drawback of requiring labor and consumables compared with leaving specimens in 96-well microplates. We predict that this investment is in most cases justified because nowadays barcoding rarely serves to only identify common species. Increasingly, barcoding is used to discover and study neglected dark taxa which requires access to vouchers for additional DNA sequencing, imaging and trait extraction (Srivathsan et al., 2023; Caruso et al., 2024; Hartop et al., 2024; Hébert & Favret, 2025; Meier et al., 2025a). The workflow is therefore timely as ecology and evolutionary biology move beyond documenting occurrence and abundance with DNA barcodes towards explaining patterns through analysis of traits and biomass (Shirali et al., 2026) as well as genomes and interactions (Brown et al., 2025). Converting barcode output into reusable collections thus allows anonymous mOTUs to accumulate biological information and develop into biological knowledge.

## Acknowledgements

This work was supported by the European Union’s Horizon Europe Research and Innovation Programme under Grant Agreement No. 101134200 “Forest surveillance with artificial intelligence and digital technologies – FORSAID”. We would also like to thank the lab technicians and students of the Center for Integrative Biodiversity Discovery for their help and assistance.

## Data Availability

All data and files necessary to reproduce the results presented in this study are provided in the Supplementary Material associated with this article. *MOTUmapper* is available at https://github.com/asrivathsan/MOTUmapper.

## Author Contributions

R.M., D.O. and V.F. conceived the ideas and designed methodology; A.S. developed MOTUmapper; H.-M.L. developed the SORTER hardware; H.W. developed the SOFTER software; V.F., L.L. and R.P. collected and analysed the data; V.F. and R.M. led the writing of the manuscript. All authors contributed critically to the drafts and gave final approval for publication.

