## Supplementary Material for "Beyond DNA barcodes: an open-source workflow for recovering and organizing barcoded vouchers for ecological and evolutionary research"

The following links point to the Figshare repositories containing the files needed to replicate this study.

All files associated with the Inner-Outer system and the “Popsicle” device (3D print files, laser engraver input, instructions): <https://doi.org/10.6084/m9.figshare.33176285>

All files associated with the SORTER platform (3D print files, hardware components, software): <https://doi.org/10.6084/m9.figshare.33174449>

Test data used in evaluation of voucher recovery time: <https://doi.org/10.6084/m9.figshare.33162197>
